# Targeted single cell expression profiling identifies integrators of sleep and metabolic state

**DOI:** 10.1101/2024.09.25.614841

**Authors:** Meng-Fu Maxwell Shih, Jiwei Zhang, Elizabeth B. Brown, Joshua Dubnau, Alex C. Keene

## Abstract

Animals modulate sleep in accordance with their internal and external environments. Metabolic cues are particularly potent regulators of sleep, allowing animals to alter their sleep timing and amount depending on food availability and foraging duration. The fruit fly, *Drosophila melanogaster*, suppresses sleep in response to acute food deprivation, presumably to forage for food. This process is dependent on a single pair of Lateral Horn Leucokinin (LHLK) neurons, that secrete the neuropeptide Leucokinin. These neurons signal to insulin producing cells and suppress sleep under periods of starvation. The identification of individual neurons that modulate sleep-metabolism interactions provides the opportunity to examine the cellular changes associated with sleep modulation. Here, we use single-cell sequencing of LHLK neurons to examine the transcriptional responses to starvation. We validate that a Patch-seq approach selectively isolates RNA from individual LHLK neurons. Single-cell CEL-Seq comparisons of LHLK neurons between fed and 24-hr starved flies identified 24 genes that are differentially expressed in accordance with starvation state. In total, 12 upregulated genes and 12 downregulated genes were identified. Gene-ontology analysis showed an enrichment for *Attacins*, a family of anti-microbial peptides, along with several transcripts with diverse roles in regulating cellular function. Targeted knockdown of differentially expressed genes identified multiple genes that function within LHLK neurons to regulate sleep-metabolism interactions. Functionally validated genes include an essential role for the E3 ubiquitin Ligase *insomniac,* the sorbitol dehydrogenase *Sodh1,* as well as *AttacinC* and *AttacinB* in starvation-induced sleep suppression. Taken together, these findings provide a pipeline for identifying novel regulators of sleep-metabolism interactions within individual neurons.

## Introduction

Dysregulation of sleep and feeding has severe health consequences. Acute sleep loss results in increased appetite and insulin insensitivity, and chronically sleep-deprived individuals are more likely to develop obesity, metabolic syndrome, type II diabetes, and cardiovascular disease (Chaput et al., 2007; Peppard et al., 2000; Taheri et al., 2004; Van Cauter & Knutson, 2008). Conversely, metabolic state potently modulates sleep and circadian behavior (Arble et al., 2015; Froy & Miskin, 2010; Laposky et al., 2008). Flies, rodents, and humans all suppress sleep or experience diminished sleep quality when starved, presumably to increase foraging (Danguir & Nicolaidis, 1979; Horne, 2009; Keene et al., 2010; Macfadyen et al., 1973; Thimgan et al., 2010). While genes and neurons have been identified that regulate both sleep and feeding behavior, much less is known about the genes and neural circuits that regulate sleep in accordance with feeding state. Given the growing epidemiological and functional evidence for sleep-feeding interactions, it is critical to understand how neural circuits modulating sleep are impacted by metabolic state.

*Drosophila* is a leading model for studying genetic regulation of sleep and metabolism, and mechanisms underlying each of these processes are highly conserved from flies to mammals (Allada & Siegel, 2008; Padmanabha & Baker, 2014; Sehgal & Mignot, 2011; Taghert & Nitabach, 2012). In addition, many genes and signaling molecules required for the integration of sleep and feeding are conserved across phyla, including those that regulate the circadian clock, energy stores, and neuropeptide regulation (Chung et al., 2017; Keene et al., 2010; Yurgel et al., 2019a). Flies, like mammals, suppress sleep when starved (Danguir & Nicolaidis, 1979; Horne, 2009; M. Yurgel et al., 2014). Because *Drosophila* only live for 2-3 days, when starved, it is likely evolutionarily adaptive to suppress sleep and increase foraging in response to food deprivation. Large-scale RNA interference (RNAi) screens have identified conserved genes that regulate behavioral and metabolic processes (Dietzl et al., 2007; Neely et al., 2010; Pospisilik et al., 2010). For example, neuron-specific RNAi screen have identified multiple genes that are required for starvation-induced sleep suppression, including the mRNA/DNA-binding protein Translin (encoded by *trsn)* and the neuropeptide Leucokinin (Murakami et al., 2016a; Yurgel et al., 2019a). Targeted approaches have identified brain nutrient sensors, insulin signaling, and the serine metabolism pathway (Brown et al., 2020; Oh & Suh, 2023; Sonn et al., 2018). These findings suggest distinct neural processes regulate changes in sleep under standard fed vs starvation conditions.

Sleep and feeding circuits are distributed throughout the brain. Multiple neurons in the brain have been implicated in both the regulation of sleep and metabolic function including the Insulin Producing Cells and the Lateral Horn Leucokinin (LHLK) neurons (Brown et al., 2019; Cong et al., 2015; Metaxakis et al., 2014; M. E. Yurgel et al., 2019a). The LHLK neurons are proposed integrators of sleep and metabolic state because they are activated during starvation, and their activity is essential for starvation-induced sleep suppression (Liu et al., 2015; M. E. Yurgel et al., 2019b; Zandawala et al., 2018). These neurons express the RNA binding protein *translin* and the neuropeptide *Leucokinin,* both of which are required for starvation-induced sleep suppression, but are dispensable for normal sleep under fed conditions (Murakami et al., 2016a). The LHLK neurons ramify throughout the central brain, with terminals that project near the mushroom bodies and insulin producing cells, two brain regions that are known to regulate sleep (Liu et al., 2015). Functional analysis suggests the Insulin-Producing Cells are targets of LHLK neurons, and that silencing of LHLK signaling in these neurons disrupts starvation-induced sleep suppression (M. E. Yurgel et al., 2019a). The identification of LHLK neurons as critical regulators of sleep– metabolism interactions provides the opportunity to apply single-cell genomics to study how feeding state regulates gene expression.

We have established a robust pipeline for transcriptional profiling of individual identified neurons and then used this approach to identify transcripts that are differentially expressed in LHLK neurons in fed vs starved animals. We then used LHLK-neuron specific RNAi-mediated to test the behavioral effects of knocking down each differentially expressed transcript. Our findings identify genes that function within LHLK neurons to mediate the effects of starvation on sleep.

## Results

The identification of LHLK neurons as critical regulators of sleep–metabolism interactions provides the opportunity to apply single-cell genomics to study how feeding state regulates gene expression. We developed a Patch-seq platform to enable single-cell RNA sequencing of neurons harvested from living animals (Fig. 1A-B and Fig. 1D). To accomplish this, we use a glass electrode to patch onto a single neuron, guided by GFP expression. To access individual LHLK neurons, we opened a window of dorsal head cuticle to visualize GFP in the *LK-GAL4>UAS-GFP* flies. The LHLK neurons were readily identifiable based on size, location and GFP expression. Once isolated from the head, we used CEL-Seq2, a linear amplification library preparation method (Hashimshony et al., 2012; Yanai & Hashimshony, 2019) that selectively enriches for a segment at the 3’ untranslated region. This strategy offers several key advantages over high-throughput single cell sequencing methods for brain-wide profiling of many thousands of neurons. First, the Patch-seq approach permits repeated targeting of the same neuron across multiple animals and even of the same neuron on both sides of one animal’s brain. Second, the linear amplification used does an excellent job of preserving the relative levels of each transcript. Finally, the CEL-Seq2 amplification approach reduces the library complexity to a defined portion of each transcript, providing highly robust detection of even relatively low abundance transcripts.

**Figure 1.**
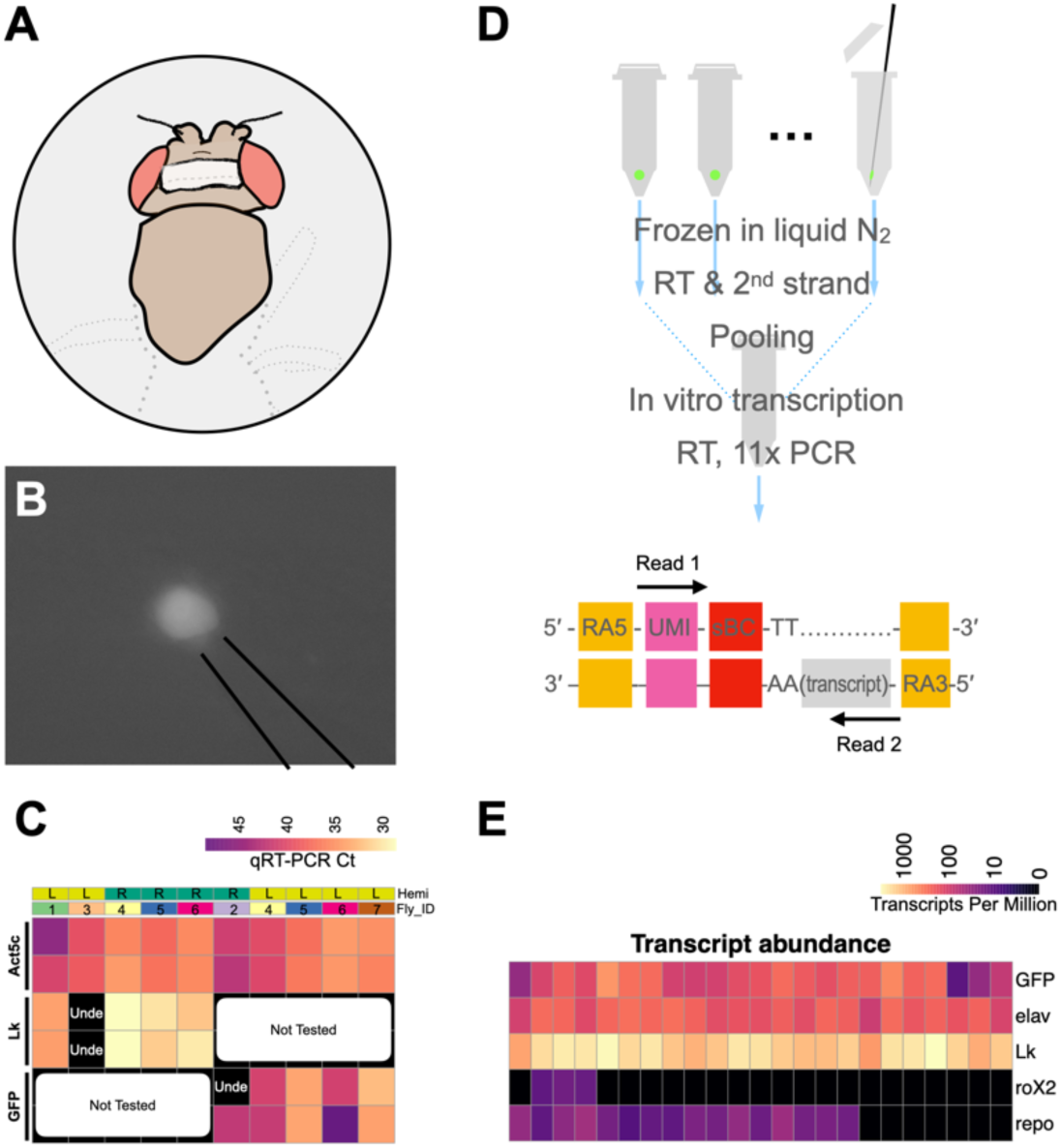
Single-cell transcriptome analysis of LHLK neurons via Patch-Seq. (**A**) A fly tethered on a custom fly stage underwent microdissection to have a window of dorsal head cuticle removed (grey shadow) and then was mounted on a rig. (**B**) Fluorescence imaging through the cuticle window allowed visualization of the GFP labeled LHLK neurons, which were then sucked into the glass electrode. (**C**) Quantitative reserve transcription PCR (qRT-PCR) was used to measure cycle threshold (Ct) values for *Lk* or *GFP*, together with the housekeeping gene *Act5c* in 10 single LHLK neurons. The control gene *Act5c* is always detected across all the single LHLK neurons bilaterally (left (L) or right (R) hemisphere (Hemi)) from seven flies (Fly_ID) and their technical replicates. *Lk* gene is reliably detected except for being undetermined (Unde) in one sample or two technical replicates. GFP is reliably detected except for in one *GFP* technical replicate where it was undetermined. (**D**) CEL-Seq2 protocol for library construction and sequencing allows routine detection of ∼6000 unique transcripts per LHLK neuron. (**E**) Fidelity of CEL-seq shown for marker gene expression from 23 different LHLK neurons. High (bright colors) expression seen for *GFP*, *Lk*, and the *elav* neuronal marker. By contrast, we do not detect the male-specific gene *roX2* (mean 2.67 TPM; median 0 TPM) or detect weak signal of the glial *repo* marker (mean 24.67 TPM; median 22.47 TPM).

We first sought to validate the specificity of the method to detect transcripts that derive from the target neurons. We extracted 10 LHLK neurons from fed flies and measured known markers of gene expression through quantitative real-time PCR. The housekeeping gene *Act5c* was detected in all samples, confirming our ability to extract intact RNA from the isolated cells. (Fig 1C; Supplemental Table 1). We also consistently detect transcripts that are expected markers of the LHLK neurons including the transgenic GFP and the neuropeptide gene *Lk*. Overall, these findings validate this approach to isolate single LHLK neurons.

We next used CEL-Seq to directly compare the transcriptional profiles of fed and starved flies carrying the same transgenes. LHLK neurons were bilaterally harvested between Zeitgeber Time (ZT) 1 and ZT5 hours, from female flies that were either fed under standard growth conditions or starved for 24 hours prior to cell harvesting (Fig. 1D; see methods). The tips of glass electrodes containing harvested individual LHLK neurons were broken into microcentrifuge tubes and flash frozen in liquid nitrogen. After single-cell reverse transcription with barcoded primers, a mix of LHLK neurons from fed and starved flies were pooled into libraries that were amplified by *in vitro* transcription and limited cycles of PCR and later sequenced on Illumina HiSeq 2500 sequencer (adapted CEL-Seq2 protocol in Supplementary File 1; Fig 2A) (Yanai & Hashimshony, 2019). A custom pipeline (see methods) was used to analyze the sequencing data and reconstruct single LHLK transcriptomes (Supplementary Table 2 & 3). Reconstructed single LHLK transcriptomes from fed flies showed expected signal for marker genes (Fig. 1E): strong signal for transgenic *GFP*, neuronal marker *elav*, and the neuropeptide gene *Lk* while the male-specific gene *roX2* and glial marker gene *repo* (Xiong et al., 1994) were barely detectable. This method also provides reliable detection across a wide range of expression levels. We routinely detected ∼6000 unique transcripts per cell. We compared the transcriptional profiles of LHLK neurons in fed and 24-hr starved flies (18 starved vs 23 fed) to identify transcripts that were differentially expressed. We identified 12 that are transcriptionally upregulated and 12 that are downregulated in accordance with feeding state. A number of genes with known roles in feeding or sleep behaviors were identified, including satiety peptide female-specific independent of transformer (*fit*) and the sleep promoting ubiquitin ligase adaptor protein *insomniac* (Q. Li et al., 2017; Pfeiffenberger & Allada, 2012; Stavropoulos & Young, 2011; Sun et al., 2017)(Figure 2B-C and Table 1 and Supplementary Table 4). Gene Ontology (GO) enrichment identified ‘anti-microbial peptides’ and ‘Attacins’, a class of anti-microbial peptide, as enriched among differentially expressed genes (Fig 2D; Supplementary Table 5 & 6)(Gene Ontology Consortium et al., 2023). Together, these differentially expressed genes provide candidate regulators of sleep-metabolism interactions.

**Figure 2.**
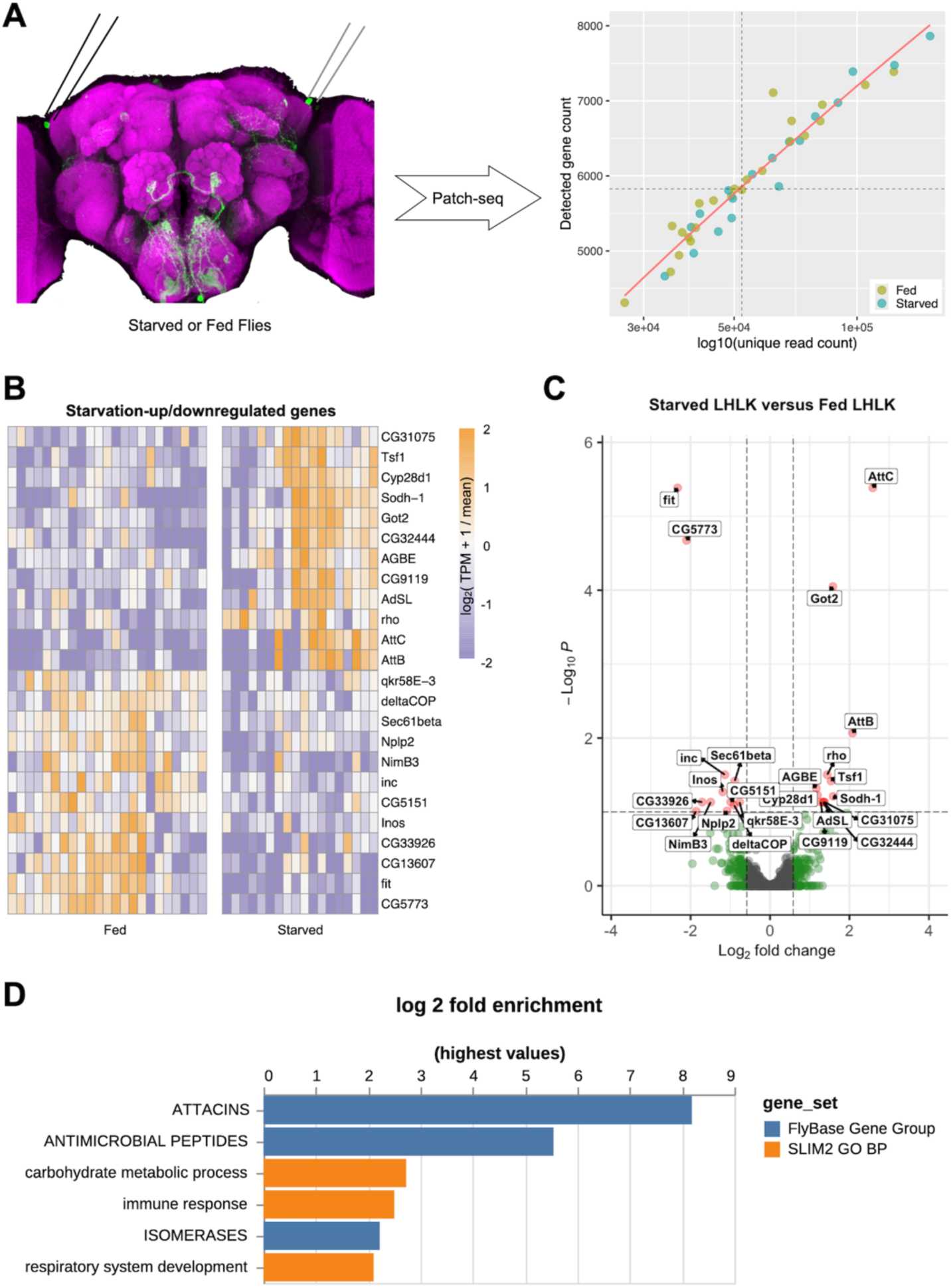
Transcriptional analysis of LHLK neurons reveals differentially expressed genes. (**A**) 24-hr of starvation (green) vs feeding-control (yellow) was followed by Patch-Seq profiling. We generated the 2000 most variably expressed genes in the differential expression analysis. Genes with an adjusted P value smaller than 0.1 are highlighted in red and labeled; genes with a fold change bigger than 1.5 are highlighted in green. (**D**) GO analysis using PANGEA. 24 genes with a DE adjusted P value less than 0.1 were used for analysis against the gene ontology subset SLIM2 GO BP (orange) and the gene group collection FlyBase Gene Group (dark blue). P-value filter of 0.1 was applied. Bar plot represents the log2 fold change of each gene set.

**Table 1.** Preliminary sequencing hits. Genes that are up or down regulated (Log2FC) in LHLK neurons following 24 hours food deprivation.

| Gene | Log<sub>2</sub>FoldChange | padj | Gene function |
| --- | --- | --- | --- |
| ***AttC* or *AttacinC*** | 2.59 | 4.1E-06 | encodes an immune inducible peptide that shows homology to antibacterial peptides |
| ***Got2*** | 1.58 | 8.9E-05 | encodes an L-aspartate:2-oxoglutarate aminotransferase involved in glutamate biosynthesis |
| ***AttB* or *AttacinB*** | 2.08 | 8.6E-03 | encodes an immune inducible peptide that shows homology to antibacterial peptides |
| ***rho*** | 1.44 | 3.1E-02 | encodes an intra-membrane serine protease that processes the membrane precursors of Egfr ligands. Its roles include growth regulation, cell survival and developmental patterning. |
| ***Tsf1*** | 1.53 | 3.8E-02 | encodes an iron binding protein induced during the immune response. Iron sequestration is a classical host defense mechanism to combat bacterial infection. |
| ***AGBE*** | 1.16 | 4.8E-02 | encodes a protein that is inferred to be a hydrolase involved in glycogen synthesis and hypoxia, based on the function of its orthologs. It contributes to the determination of lifespan. |
| ***Cyp28d1*** | 1.19 | 5.9E-02 | May be involved in the metabolism of insect hormones and in the breakdown of synthetic insecticides. |
| ***Sodh-1* or *Sodh1*** | 1.59 | 6.2E-02 | Predicted to enable L-iditol 2-dehydrogenase activity. Involved in fructose biosynthetic process and sorbitol catabolic process. |
| ***CG32444*** | 1.35 | 7.4E-02 | Predicted to enable aldose 1-epimerase activity. Predicted to be involved in galactose catabolic process via UDP-galactose and glucose metabolic process. |
| ***CG9119*** | 1.28 | 7.4E-02 | Predicted to enable hydrolase activity, acting on ester bonds and zinc ion binding activity. Predicted to be active in nucleus. |
| ***AdSL*** | 1.34 | 7.4E-02 | encodes a protein that might be a lyase involved in purine nucleotide metabolic process based on orthology. |
| ***CG31075*** | 1.37 | 7.4E-02 | Predicted to enable aldehyde dehydrogenase (NAD<sup>+</sup>) activity. Predicted to be involved in pyruvate metabolic process. Predicted to be located in mitochondrion. |
| ***fit*** | -2.33 | 4.1E-06 | NA |
| ***CG5773*** | -2.10 | 2.1E-05 | NA |
| ***inc* or *insomniac*** | -1.14 | 3.1E-02 | encodes a BTB domain-containing protein that is essential in neurons for normal sleep-wake cycles. |
| ***Sec61beta* or *Sec61β*** | -0.89 | 3.8E-02 | encodes a protein involved in the regulation of autophagy. |
| ***Inos*** | -1.19 | 5.4E-02 | Key enzyme in myo-inositol biosynthesis pathway that catalyzes the conversion of glucose 6-phosphate to 1-myo-inositol 1-phosphate in a NAD-dependent manner |
| ***CG33926*** | -1.71 | 7.4E-02 | Involved in defense response to virus |
| ***NimB3*** | -1.49 | 7.4E-02 | Predicted to be located in extracellular region |
| ***qkr58E-3*** | -0.77 | 7.4E-02 | encodes an RNA binding protein that contributes to mRNA splicing and apoptosis. |
| ***CG5151*** | -0.99 | 7.4E-02 | NA |
| ***DeltaCOP* or *δCOP*** | -0.90 | 8.1E-02 | mutants show narrow tracheal tubes due to defective luminal extracellular matrix secretion. |
| ***Nplp2*** | -1.08 | 9.7E-02 | Neuropeptide gene involved in humoral immune response. |
| ***CG13607*** | -1.86 | 9.8E-02 | NA |

We sought to investigate the role of genes that are transcriptionally regulated by feeding state on sleep regulation. We genetically expressed RNAi lines targeted to identified differentially expressed genes within LHLK neurons and measured sleep under fed and starved states. Briefly, flies were placed in *Drosophila* Activity Monitors (DAMs), and sleep was measured for 24 hours on food, followed by 24 hours on agar (starved condition). We first tested genes that were upregulated after 24 hours of starvation. Knockdown of each of 10 such candidate genes did not impact sleep during the fed state for any of the lines tested (Fig 3, Fig S1 and Table 2). Similarly, no differences in waking activity were observed in fed flies for any of the lines tested (Supplemental Table 8). However, flies with knockdown of (*sorbitol dehydrogenase 1*) *Sodh1* in LHLK neurons slept significantly more than controls when starved, suggesting upregulation of *Sodh1* is required for starvation-induced sleep suppression (Fig 3A,B). Knockdown of *Sodh1* also resulted in reduced waking activity during the starved state, but not in fed flies, fortifying the notion that *Sodh1* selectively regulates starvation-dependent changes in sleep (Fig S2A). We used a Markov model that predicts sleep and wake drive, providing indicators of sleep quality (Wiggin et al., 2020). Wake drive was reduced, while sleep drive was not significantly altered but trended towards significance (Fig S2B,C). Similarly, knockdown of *Attacin B* (*AttB)* or *Attacin C* (*AttC*) in LHLK neurons increased sleep in starved, but not fed, flies (Fig 3C-F). Knockdown of *AttB* or *AttC* led to reduced waking activity under starvation conditions, as well as reduced wake drive and increased sleep drive, without impacting these variables under fed conditions (Figure S2D-I). Together, these findings support the notion that elevated levels of *Sodh1*, *AttB* and *AttC* are required for starvation-induced sleep suppression.

**Figure 3.**
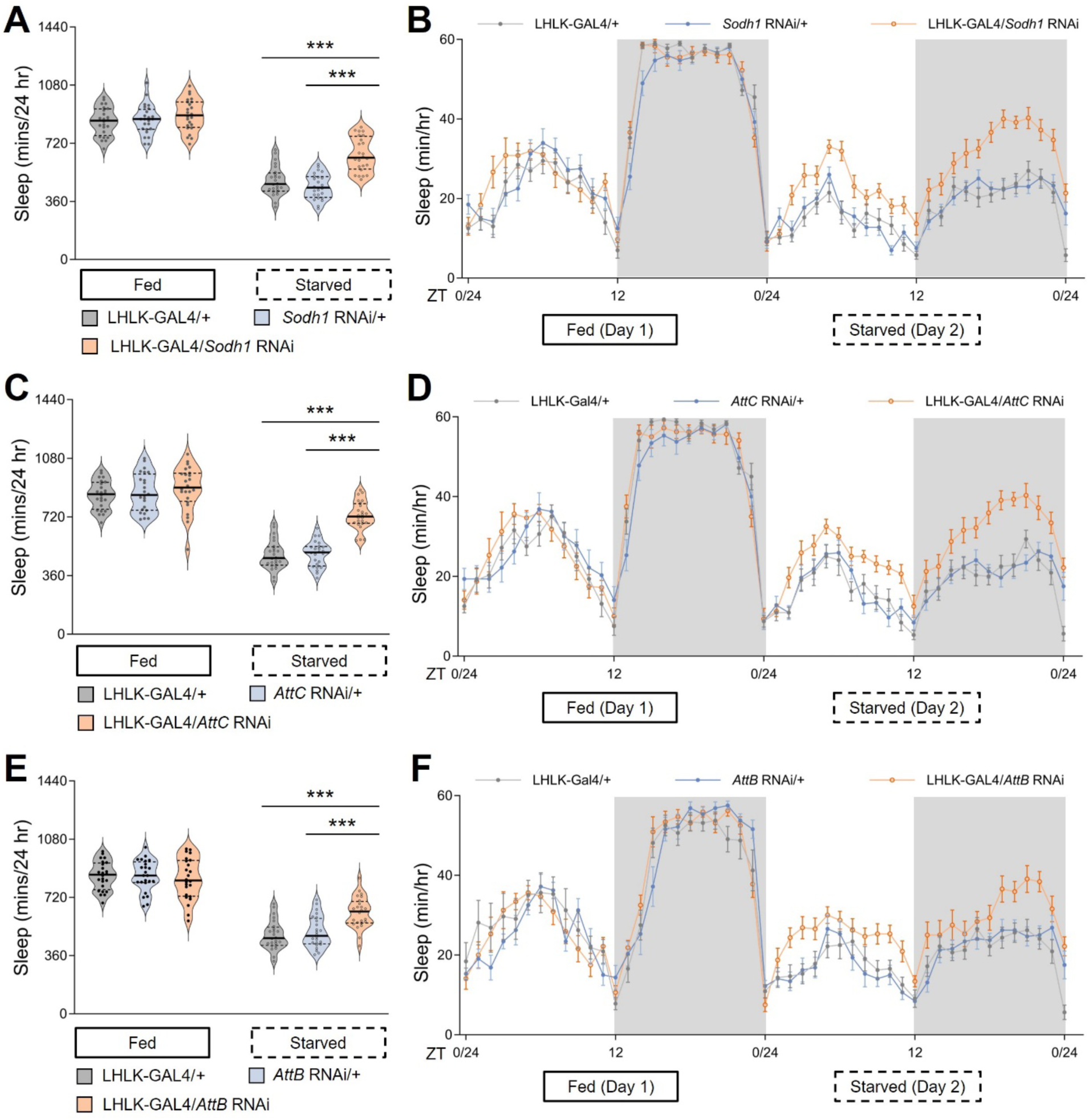
Transcriptional reduction of Sodh1 or Attacins in LHLK neurons mitigates starvation-induced sleep suppression. **(A)** Total sleep duration reveals that sleep in LHLK-GAL4>*Sodh1*^RNAi^ flies (black dots in light apricot shadow) does not differ from control flies using LHLK-GAL4 (dark cycles in light gray shadow) or RNAi line (dark cycles in light blue shadow) crossed to the wild-type genetic background (+) respectively under fed conditions (Day 1, P = 0.2758), but is increased during the starved state (Day 2). Two-way repeated-measures ANOVA: F _(2, 138)_ = 10.49, p < 0.0001 and Tukey’s multiple comparison tests: P < 0.0001, Kruskal-Wallis test for three groups following food deprivation, P < 0.001. All columns mean ± SEM. N = 22-25 each group. Legend key applies to A-F. (**B**) Sleep profile of hourly sleep averages over a 48 hr experiment under fed and stared conditions. Gray box represents night phase. ZT denotes Zeitgeber time, where ZT 0-12 corresponds to lights-on, and ZT 12-24 lights off. Sleep is reduced on starvation day for all genotypes, but sleep suppression by food deprivation is alleviated in mutant fruit flies expressing *Sohd1*-RNAi in the LHLK neurons (light apricot) compared to GAL4 and RNAi controls (light gray and light blue respectively). (**C**) Total sleep duration reveals that sleep in LHLK-GAL4>*AttC*^RNAi^ flies does not differ from controls under fed conditions (P = 0.3448), but is increased during the starved state. Two-way repeated-measures ANOVA: F _(2, 138)_ = 16.54, p < 0.0001 and Tukey’s multiple comparison tests: P < 0.0001. Kruskal-Wallis test for groups under starvation, P < 0.001. (**D**) Sleep profile reveals that sleep in LHLK-GAL4>*AttC*^RNAi^ flies (light apricot) and control flies under fed condition and the starved state. (**E**) Total sleep duration shows no significant differerence between LHLK-GAL4>*AttB*^RNAi^ flies and controls under fed conditions (P = 0.7788), but sleep is induced during the starved state in mutant flies compared to two control groups. Two-way repeated-measures ANOVA: F _(2, 138)_ = 8.908, p < 0.0001 with and Tukey’s multiple comparison tests: P = 0.0082. Kruskal-Wallis test, P < 0.001. (**F**) Daily sleep profile of hourly sleep averages in LHLK-GAL4>*AttB*^RNAi^ flies compared to controls. All data are mean ± SEM; *P < 0.05; **P < 0.01; ***P < 0.001.

Next, we tested genes that were downregulated within LHLK neurons in response to starvation. Knockdown of *insomniac* (*inc*) and *CG5151* significantly reduced sleep in the fed state, while none of the other genes tested reduced sleep in the fed state (Fig 4A,B). In addition, knockdown of *inc* increased sleep in starved flies (Fig. 4A). These findings are consistent with the notion that downregulation of *inc* is required for metabolic regulation of sleep in flies, while *CG5151* selectively regulates sleep during the fed state. Only knockdown of *Sec61β* increased sleep in fed flies, and sleep was also increased in starved flies (Fig 4C). Sleep in starved conditions, but not fed conditions, was increased in flies with LHLK-specific knock-down of *δCOP*, *CG5773*, or *fit* (Fig 4D-F). This phenotype reveals complex interactions between gene expression and sleep phenotype. While being differentially expressed predicts functional impact on modulation of sleep by feeding state, the sign of the effect on transcript levels does a poor job of predicting the direction of effect on modulation of sleep by starvation. This may reflect the fact that we measured changes in transcript abundance at a single timepoint after the onset of starvation. If the transcriptional changes over time are dynamic, then the sign of effect from chronic knock down may be hard to predict.

**Figure 4:**
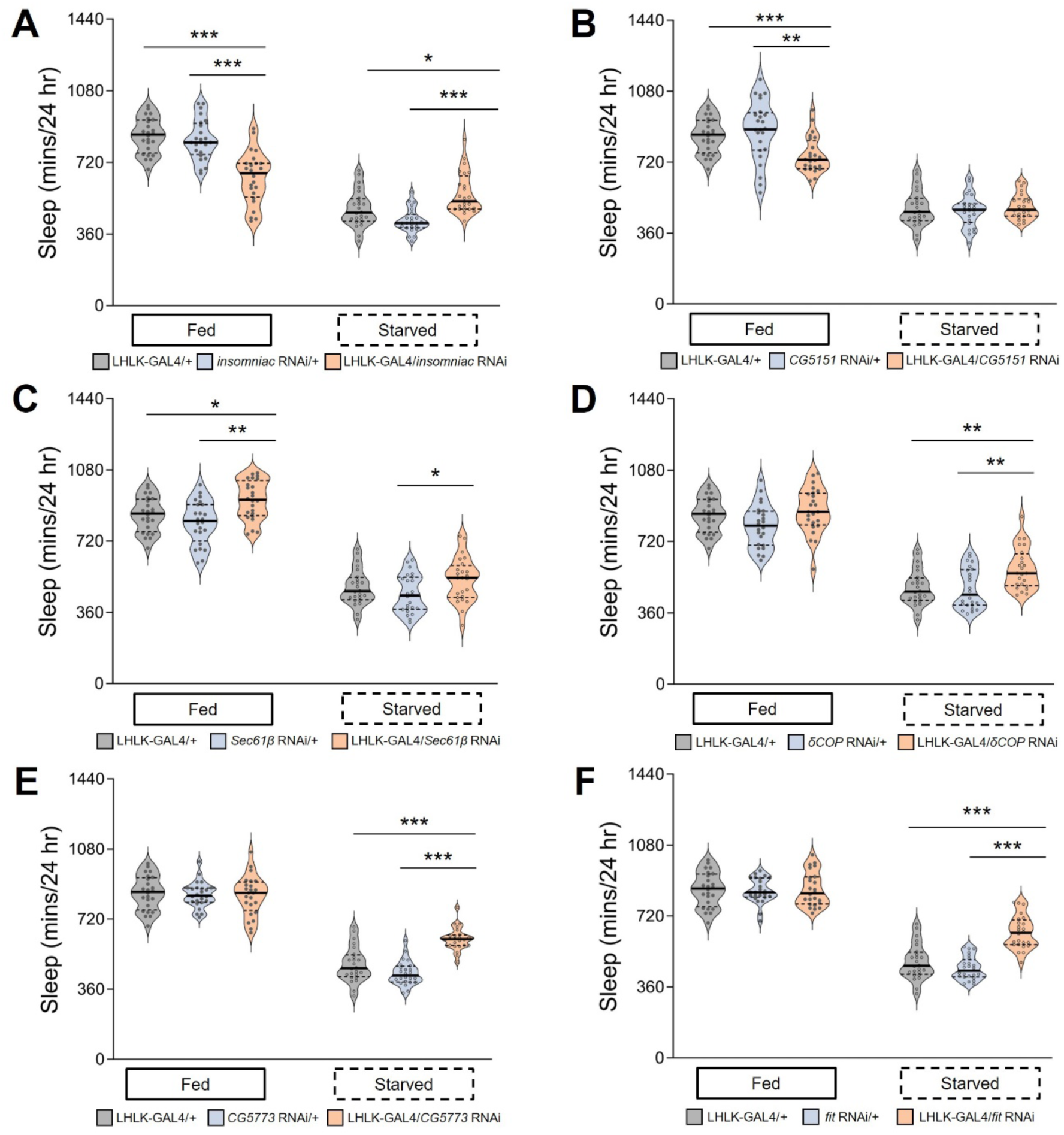
Behavioral role for genes downregulated in LHLK neurons following food deprivation. For each RNAi line, sleep was measured for 24hrs on food, followed by 24hrs on agar for starvation. (**A**) Sleep duration is reduced on food (P < 0.001) and increased on agar in LHLK-GAL4>*insomniac*^RNAi^ flies (light apricot shadow) compared to LHLK-GAL4 controls (light gray shadow, P= 0.0253) or RNAi controls (light blue shadow, P < 0.001), which crossed to the wild-type genetic background (+) respectively. Two-way repeated-measures ANOVA: F _(2, 138)_ = 40.00, p < 0.0001. Krusal-Wallis test for three groups under fed or starved conditions, P < 0.001 All columns mean ± SEM. N = 22-25 each group. Legend key applies to **A**-**F**. (**B**) Daily total sleep is only decreased under fed condition (P = 0.0084) no significant change in starved state (P = 0.7653) for *CG5151* knockdown flies compared to controls. Two-way repeated-measures ANOVA: F _(2, 138)_ = 7.016, p < 0.0013. Krusal-Wallis test for three groups under fed condition with unpaired t test for P value correction. (**C**) Sleep was significantly increased in both fed or starved states (P =0.0019, P = 0.0206 respectively) in LHLK-GAL4>*Sec61β*^RNAi^ flies compared with the corresponding controls under the same situation. Two-way repeated-measures ANOVA: F _(2, 138)_ = 66.73, p < 0.0001. Krusal-Wallis test. **D**-**F**) Sleep duration is normal on food, but significantly increased on agar compared to control flies in LHLK-GAL4>*δCOP*^RNAi^ flies (**D**, two-way ANOVA, F _(2, 138)_ = 1.715, p < 0.0001, and P = 0.0020, Kruskal-Wallis test for starved flies), LHLK-GAL4> *CG5773*^RNAi^ flies (**E**, two-way ANOVA, F _(2, 138)_ = 17.33, p < 0.0001, and P < 0.001), and LHLK-GAL4>*fit*^RNAi^ flies (**F**, two-way ANOVA, F _(2, 138)_ = 17.17, p < 0.0001, and P < 0.001). Krusal-Wallis test for three groups and unpaired t test for comparison of two groups. Data are mean ± SEM. *P < 0.05; **P < 0.01; ***P < 0.001.

To further investigate the impact of each gene on metabolic regulation of sleep, we calculated the effects of each knock down on the total percentage of sleep loss of individual flies during the starvation state. Over 24 hours of starvation, wild-type flies normally lose 30-50% of their sleep (Fig 5). Using this analysis, we found that a large fraction of the identified DE genes had palpable impact on dietary modulation of sleep. Starvation-induced sleep loss was significantly reduced in flies with knockdown of *inc*, *CG5151*, *δCOP*, *CG5773*, or *fit,* all of which are downregulated during starvation. Similarly, knockdown of *Sodh1*, *AttC*, *AttB*, *Got2*, or *Cyp28d1,* that have elevated expression during starvation, results in reduced magnitude of dietary modulation of sleep. Taken together these functional studies provide evidence that feeding-associated transcriptional changes in LHLK neurons play a critical role in modulation of sleep by satiety state.

**Figure 5.**
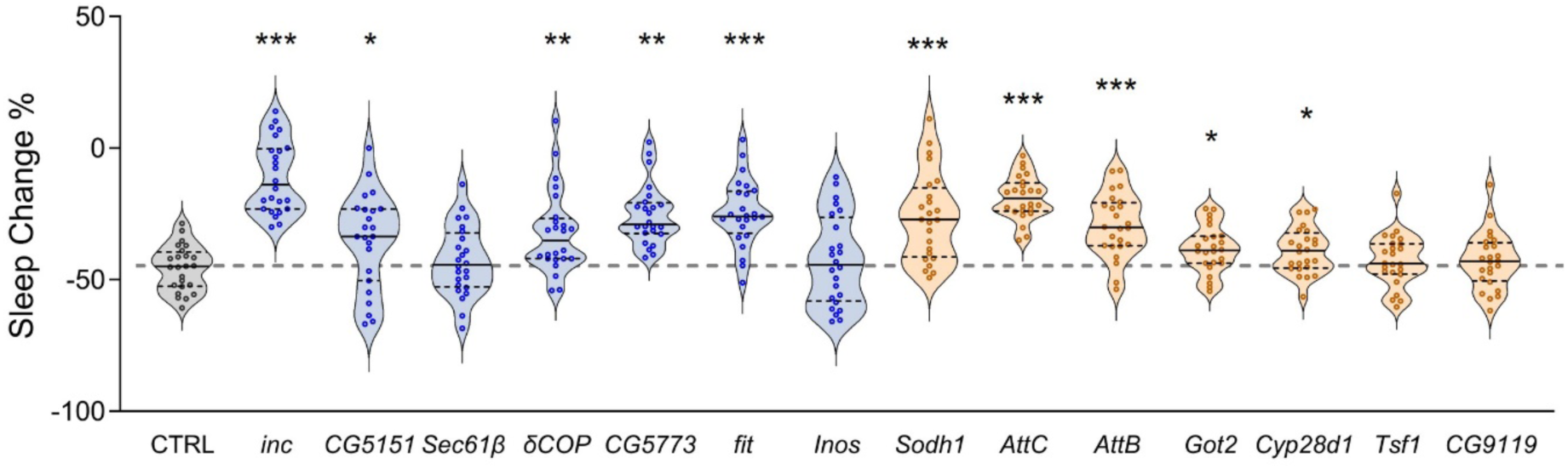
Starvation-induced sleep loss is reduced by knockdown of up-regulated or down-regulated candidate genes. Starvation-induced sleep suppression was calculated based on the percentage change in sleep between the fed and starved state. LHLK-GAL4/+ control flies (grey) suppressed sleep by 45-50% (grey dotted line). Of the transcriptionally downregulated genes (blue), starvation induced sleep loss was reduced in LHLK-GAL4>*inc*^RNAi^ (P < 0.001), LHLK-GAL4>*CG5151*^RNAi^ (P = 0.0142), LHLK-GAL4>*δCOP*^RNAi^ (P = 0.0070), LHLK-GAL4>*CG5773*^RNAi^ (P < 0.001) and LHLK-GAL4>*fit*^RNAi^ (P < 0.001) flies. Of the transcriptionally up-regulated genes (apricot), starvation-induced sleep suppression was reduced in LHLK-GAL4> *Sodh1*^RNAi^ (P < 0.001), LHLK-GAL4>*AttC*^RNAi^ (P < 0.001), LHLK-GAL4>*AttB*^RNAi^ (P < 0.001), LHLK-GAL4>*Got2*^RNAi^ (P = 0.0063), and LHLK-GAL4>*Cyp28d1*^RNAi^ (P = 0.0101) flies. One-way ANOVA: F _(14, 344)_ = 15.50, p<0.0001 followed by Brown-Forsythe test, P = 0.0005. N = 24-25 per group. Data are mean ± SEM; *P < 0.05; **P < 0.01; ***P < 0.001.

## Discussion

Sleep is governed by both internal state and external stimuli. Understanding the neural and genetic basis for these interactions is a central challenge in sleep research. In *Drosophila*, numerous neuronal cell types and brain regions have been implicated in sleep regulation including circadian neurons, the mushroom bodies, the fan-shaped body, and catecholamine neurons. The LHLK neurons are unique for their role in regulating sleep in a context-specific fashion. Silencing LHLK neurons has no effect on sleep in fed flies, but impairs starvation-induced sleep suppression (Yurgel et al., 2019a). In the starved state, the LHLK neurons become more active, suggesting that transcriptional or neural circuit changes drive state-specific differences in LHLK physiology. In addition, these neurons are responsive to changes in osmolality and are critical for the formation of water memories (Senapati et al., 2019). These findings suggest LHLK neurons are regulated by nutrient availability and have a critical role in state-dependent modulation of behavior. Here, we have used state-specific single cell sequencing, followed by systematic genetic knock-down to functionally validate roles of differentially expressed genes to identify context-specific sleep regulators that act within LHLK neurons to modulate sleep.

The application of single cell sequencing approaches has been broadly used to define cell types in *Drosophila* and other model organisms (Senapati et al., 2019). More recently, these approaches have been used to examine transcriptional changes that occur across each of the cell types of the brain including during aging, Alzheimer’s-like progression, and sleep loss (H. Li et al., 2022; Lu et al., 2023; Park et al., 2024). These studies have identified transcriptional changes that occur across cell types in the brain, or the full body. Such high throughput single cell sequencing approaches offer the advantage of providing an unbiased survey of transcriptional profiles across the cell types in the brain, but they typically have poor sensitivity to detect low abundance transcripts, and do not reliably profile individual identified neurons or neuronal cell types that exist in small numbers per brain. For many *Drosophila* behaviors, including courtship, memory, circadian behavior, and sleep, the underlying circuits can include functionally relevant neuronal cell types that exist in small numbers, or even as individual identified neurons (Lee & Wu, 2020; Ly et al., 2018; Nässel & Zandawala, 2019; Shafer & Keene, 2021). A number of approaches have been used for targeted RNA seq of defined populations of behaviorally-relevant neurons. For example, fluorescence-activated cell sorting (FACS) followed by transcriptomic analysis has been used to isolate and sequence ∼200 neurons that regulate circadian behavior (Shih et al, 2019). We used a sequencing approach that is uniquely suited to transcriptionally-profile individual neurons that control behavior. The Patch-seq approach we have applied also could be used to transcriptionally profile single neurons that have been implicated in feeding, courtship, and memory (Haynes et al., 2015; Kimura et al., 2008; Yapici et al., 2016).

To probe the unique functional roles that the LHLK neurons play in the modulation of sleep by satiety, we developed a targeted single cell sequencing platform to profile expression specifically in these individual neurons, isolated bilaterally from living animals. Once isolated by patching individual LHLK neurons and physically extracting them from the brain, we profiled expression in each cell via CEL-seq, a method that provides excellent depth of sequencing, enabling detection of approximately 6000 unique transcripts per LHLK neuron. Other studies have used patch-seq to perform both electrophysiological and transcriptomic characterization of single cells (Cadwell et al., 2017). This approach has been used for mammalian cells in slice culture, as well as *in vivo* recordings, and neurons from embryonic brains (Zeisel et al., 2015). While the small size of *Drosophila* neurons makes this approach technically challenging, CEL-seq has been used to characterize motoneurons, and sleep-regulating mushroom body neurons (Crocker et al., 2016; Jetti et al., 2023). The small size of LHLK neurons made it difficult for us to perform patch-clamp physiology followed by extraction, which would have allowed us to correlate starvation-mediated increased activity of these neurons with gene expression. Future optimization of protocols may allow for combined patch-clamp physiology followed by transcriptomic analysis in LHLK and similar central brain neurons. But we were able to reliably extract these cells from the brain to produce CEL-seq libraries.

Using this approach, we identified transcripts that are differentially expressed in LHLK neurons in accordance with feeding state and then functionally tested whether such identified transcripts regulate starvation-induced sleep suppression. Previous studies have demonstrated that the mRNA/DNA binding protein *translin* (*trsn*) and the neuropeptide Leucokinin are required within LHLK neurons for animals to suppress sleep during starvation (Murakami et al., 2016a; M. E. Yurgel et al., 2019a). Our transcriptional analysis did not identify state-dependent changes in *trsn* or Lk, suggesting that the levels of their transcripts are not modulated within LHLK neurons according to feeding state. Alternatively, it is possible that changes in the expression of these two transcripts occur at timepoints that we did not monitor, or that changes in their expression were beyond the detection of our method. It is also possible that post-transcriptional modifications rather than transcriptional changes underlie the functional roles of these genes in the response to starvation.

RNAi-based screening to knock down expression levels of transcripts that we identified as differentially expressed identified numerous genes that function in LHLK neurons to regulate sleep. Of the 11 genes tested with RNAi, starvation-induced sleep loss is impaired in flies with LHLK-specific knockdown of *inc, CG5773*, *fit, Got2, Sodh1*, *AttC*, and *AttB*, phenocopying loss of *translin* and Leucokinin. *Sodh1* represents a particularly interesting example because, similar to *translin*, it is up reregulated during starvation, and selectively regulates sleep suppression in the starved state. The *Drosophila* genome encodes for two sorbitol dehydrogenase genes that are critical for cellular synthesis of glucose and fructose (Luque et al., 1998). Sorbitol dehydrogenase deficiency in *Drosophila* is associated with loss of mitochondrial function and neurodegeneration suggesting a critical role in the maintenance of neuronal homeostasis (Cortese et al., 2020; Zhu et al., 2023). Furthermore, humans with sorbitol dehydrogenase deficiencies have an increased propensity for diabetic neuropathy (Sekiguchi et al., 2019). While an acute role of *Sodh1* in modulation of sleep neural function has not been investigated, these findings suggest Sodh1 and sorbitol metabolism may play a role in metabolic regulation of sleep.

Both GO-term analysis and functional sleep analysis suggest a critical role for *Attacins* in starvation-induced sleep suppression. *AttB* and *AttC* are upregulated in LHLK neurons in response to starvation, and LHLK-specific knockdown of either gene results in increased sleep under starved, but not fed conditions. *Attacins* are anti-microbial peptides that have been implicated in the recognition of gram-positive bacteria (Buonocore et al., 2021). In *Drosophila*, numerous regulators of the immune system regulate sleep, including the anti-microbial peptide *nemuri* that improves survival rate following infection and is critical for sleep homeostasis (Toda et al., 2019). Therefore, it is possible that *Attacins* and other anti-microbial peptides serve to acutely regulate sleep in accordance with feeding state.

In addition, numerous transcripts that are down-regulated during starvation were identified as modulating sleep in LHLK neurons. For example, loss of the ubiquitin-ligase adaptor protein *inc* leads to a reduction of sleep in the fed and starved states, phenocopying pan-neuronal loss of *inc* (Pfeiffenberger & Allada, 2012; Stavropoulos & Young, 2011). *inc-*dependent ubiquitination is required for homeostatic plasticity, and flies deficient for *inc* have neuroanatomical defects suggesting both acute and developmental defects associated with *inc* mutation (Q. Li et al., 2017). In addition, knockdown of a number of genes in LHLK neurons resulted in loss of sleep, despite the finding that these genes were reduced in starved flies. The identification of genes that function within LHLK neurons to promote or suppress sleep raises the possibility that gene expression, and function, within LHLK neurons is highly dynamic. Future studies examining the differences in activity of LHLK neurons between the fed and the starved state could help identify the effects of these genes on physiology.

## Methods

### Drosophila husbandry

Flies were grown and maintained on standard *Drosophila* media (Bloomington Recipe, Genesee Scientific, San Diego, California) in incubators (Powers Scientific, Warminster, Pennsylvania) at 25°C on a 12:12 LD cycle with humidity set to 55–65%. The *w*^1118^ (#5905) flies were obtained from the Bloomington Stock Center. The LK-GAL4 line was generated by Young Joon-Kim and has previously been described (M. E. Yurgel et al., 2019a). The UAS-*RNAi* lines were obtained from the Vienna *Drosophila* Resource Center or the Bloomington Stock Center (Dietzl et al., 2007; Ni et al., 2009; Perkins et al., 2015). The stock numbers of all lines used for screening are described in Supplemental Table 1 unless otherwise stated. Mated females aged 3-to-5 days were used for all behavioral experiments performed in this study.

For Patch-seq profiling, 4-9 days old adult flies were selected for 24-hr starvation by replacing fly food with agar gel. *LK-GAL4* were crossed with a GFP reporter generated by Janelia Research Campus, *P{10XUAS-IVS-GFP-p10}attP2*. For both the starvation group in vials with agar gel or the control group in vials with regular fly food, we housed approximately 20 sex-mixed flies. Vials were placed on their side in the temperature- and humidity-controlled incubator.

### Single-cell harvesting

After 24-hr of feeding vs starvation, female flies were anesthetized on ice and tethered on a custom fly stage that was originally designed for *in vivo* patch-clamp recording (Murthy & Turner, 2013). Single-cell harvesting followed a detailed customized protocol (Supplementary File 1). In brief, after immersing the fly head in isotonic PBS, microdissection was conducted to remove the posterior dorsal lateral head cuticle to allow access to the LHLK neurons. A custom rig equipped with micromanipulator, infra-red DIC and fluorescent optics were used to harvest the individual LHLK neurons from both sides of the animal’s brain when possible. After harvesting each LHLK neuron, the tip of the electrode containing the contents of the cell was broken into a 0.5 mL microcentrifuge tube prefilled with PBS with RNase inhibitor. The tube was briefly mixed, flash frozen in liquid nitrogen, and stored at -80℃ before library construction. Sampling of cells from starved vs fed groups was performed in an alternating manner.

### Single-Cell Quantitative Reverse Transcription-PCR (qRT-PCR)

The same single-cell harvesting protocol was followed with minor differences (Supplementary File 1). For single-cell qRT-PCR, the glass electrodes were prefilled with filtered isontonic PBS. PCR tubes (0.2 mL) prefilled with RT mix1 solution that is comprised of 1.1 μL of 50 μM oligo(dT)_20_ (ThermoFisher) and 0.9 μL of RNasin Plus (Promega) were used to break the tip of glass electrode and store the single LHLK neurons at –80℃. After taking out of –80℃ freezer, individual tubes were incubated at 65 ℃ for 5 minutes and then immediately cooled on ice for 1 minute followed by brief centrifugation. RT mix2 solution that is comprised of 4 μL of 5x Superscript IV buffer, 1 μL of 0.1M DTT, 1 μL of 10 mM dNTP mix, 0.3 μL of RNasin Plus, 1 μL of Superscript IV, and 9.2 μL of RNase-Free water was added into each tube, which was then incubated at 50℃ for 10 minutes and then at 80 ℃ for 10 minutes. The RT product in each tube was split into 2 technical replicates for the housekeeping gene *Act5c* (TaqMan Gene Expression Assay for Act5C, Dm02371594_s1, ThermoFisher) and 2 technical replicates for the target gene (either *Lk ,*TaqMan Gene Expression Assay for Leukokinin Dm01843317_s1, or *GFP*, a custom TaqMan Gene Expression Assay) followed by the TaqMan default protocol. Quantitative PCR was run on the StepOnePlus Real-Time PCR System (SN: 272006700) with an adjustment of 50 cycles in total.

### Single-Cell RNA Sequencing and Analysis

An adapted CEL-Seq2 protocol was used for single-cell RNA sequencing library construction and sequencing (Supplementary File 1). In brief, 18 to 28 harvested LHLK neurons were processed in parallel for individual library constructions. Single-cell RT was conducted with primers that contain sequences of T7 promoter, Illumina RNA 5’ adaptor (RA5), hexamer UMI, single-cell barcode, and poly-T ending with V (not T), followed by second-strand synthesis. Then, all the samples were pooled together for *in-vitro* transcription-based amplification. A second round of RT was conducted with a primer that contain 3’-end common sequence of Illumina RNA PCR Index Primer sets and random hexamer, followed by 11-cycles of PCR. After 2 rounds of bead cleanup, the libraries were quantified by Qubit, size-profiled by Bioanalyzer, and sequenced on Illumina HiSeq 2500 with Rapid paired-end 25X50 configuration.

A custom script was used to string FASTX-Toolkit for preprocessing, UMI-tools for UMI-related steps (Smith et al., 2017), STAR for alignment (Dobin et al., 2013), htseq-count for counting the aligned reads (Anders et al., 2015), and samtools for alignment file formatting (H. Li et al., 2009). The count tables were analyzed by R libraries such as datatable, DESeq2, ggplot2, EnhancedVolcano, for differential expression and visualization (Blighe et al., 2018; Love et al., 2014).

*Drosophila* reference genome dm6 was used for alignment. We used the *directional* method in UMI-tools for UMI-based deduplication (Smith et al., 2017). We used criteria of ‘spike-in read over transcript read ratio’ smaller than 0.09, spike-in linearity or Pearson Correlation Coefficient between the spike-in molecules and the detected read counts bigger than 0.8, and at least 250,000 aligned reads to identify the good-quality transcriptomes for differential expression analysis. We used PANGEA (https://www.flyrnai.org/tools/pangea/) for GO analysis (Hu et al., 2023).

### Sleep Analysis

Flies were acclimated to experimental conditions for 24 hours prior to the start of all sleep analyses. Measurements of sleep were then measured over the course of 48 hours starting at ZT0 using the *Drosophila* Locomotor Activity Monitor (DAM) System (Trikinetics, Waltham, MA, USA), as previously described (Garbe et al., 2015; Hendricks et al., 2000; Shaw et al., 2000). For each individual fly, the DAM system measures activity by counting the number of infrared beam crossings over time. These activity data were then used to calculate sleep, defined as bouts of immobility of 5 min or more, using the *Drosophila* Sleep Counting Macro (Pfeiffenberger et al., 2010), from which sleep traits were then extracted. Waking activity was quantified as the average number of beam crossings per waking minute, as previously described [48]. After one day of measurement on food, flies were then transferred to tubes containing 1% agar, as previously described (Keene et al., 2010). Starvation-induced sleep loss was calculated as the difference in sleep between the fed state and the starved state as previously described(Keene et al., 2010).

### Statistical analysis

All data produced in RNAi screening are presented as means ± SEM and all hits have been validated twice. Unless stated otherwise, comparisons of overall sleep architectures in 24-hours between two or more genotypes with one treatment were performed using a one-way ANOVA followed by Tukey’s post-hoc test. Two-way repeated-measures ANOVA followed by Tukey’s post-hoc test was used for comparisons of sleep parameters between two or more genotypes with two treatments to evaluate interactions of food availability and genotype-dependent effects on sleep. The within fly percentage change in sleep was calculated as previously described (Keene, et al, 2010) and a Kruskal-Wallis test followed by Dunn’s post hoc was used for two or more genotypes, and non-parametric t tests were used for correcting p-values if stated. All statistical analyses were performed using InStat software GraphPad Prism (version 10.3.0 for Windows, Boston, Massachusetts USA).

### Data availability

All *Drosophila* strains are available upon request. Supplementary files are available at FigShare. File S1 contains the single-cell harvesting and RNA sequencing protocol. Table S1 contains the data of the single-cell qRT-PCR experiments. Tables S2 and S3 contain the unique read counts and normalized Transcript Per Million mapped reads (TPM), respectively. Table S4 contains the differential expression analysis for the 2,000 most variable genes expressing in LHLK neurons. Table S5 and S6 contain the Gene Sets selected for GO analysis and their results, respectively. Table S7 contains the fly strains used in the sleep behavioral screening and their results. The sequencing read files are available in NCBI GEO (GSE 272264). All code used to analyze single cell RNA-seq results and create the figures and tables in this manuscript are available upon request.

**Figure Supplemental 1.**
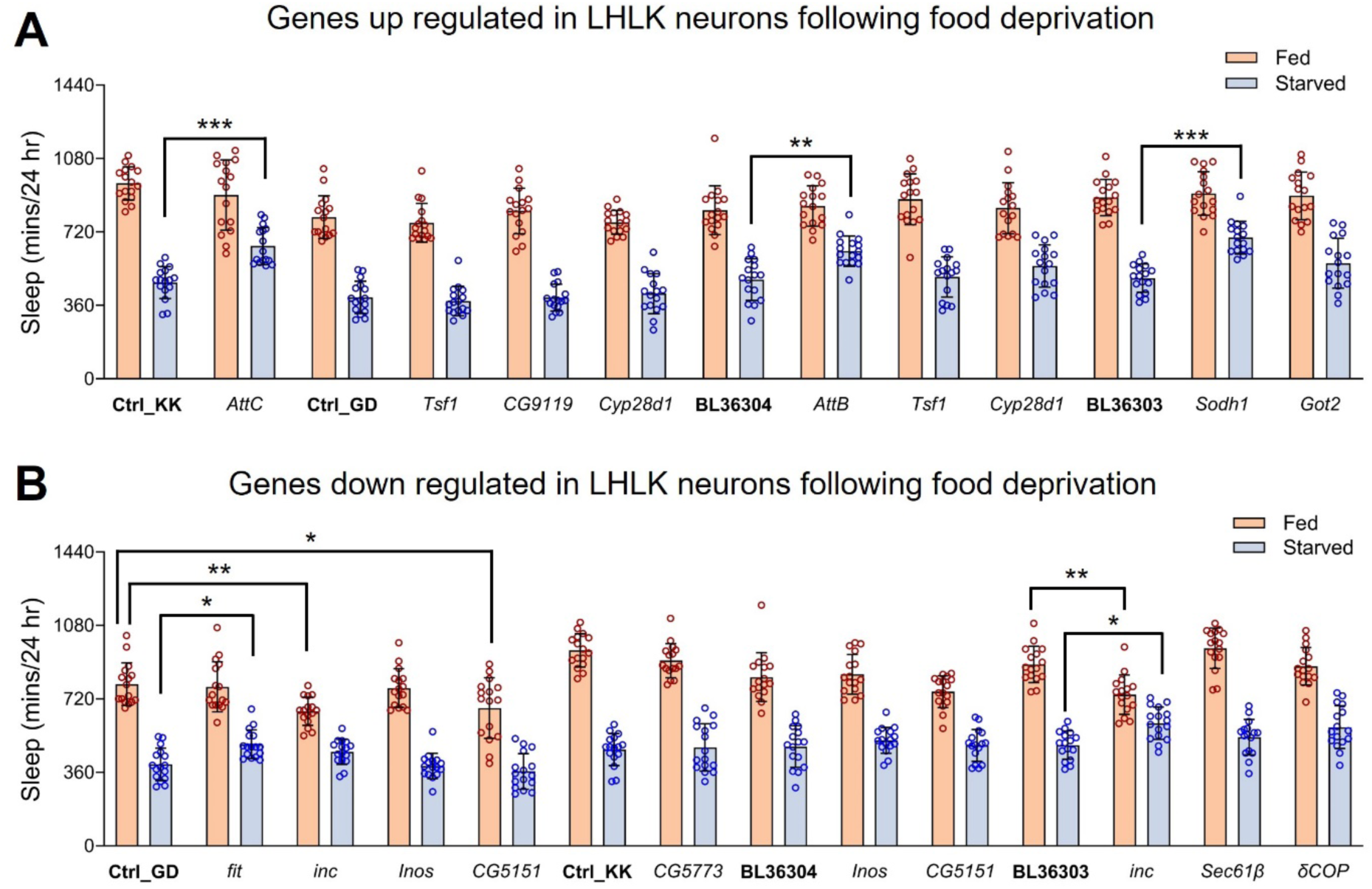
Preliminary RNAi screening for genes up-regulated (A) or down-regulated (B) in LHLK neurons following 24 hours food deprivation. (**A**) Daily sleep is less reduced in *AttC^RNAi^* (P < 0.001), *AttB^RNAi^*(P = 0.0065), or *Sodh1^RNAi^* (P<0.001) flies compared to the corresponding controls in response to 24hr starvation. Ctrl_KK or Ctrl_GD indicates control lines for RNA interference (RNAi) stocks from Vienna Drosophila Resource Center (VDRC). BL36303 and BL36304 are used as controls in screening TRiP stocks from Bloomington Drosophila Stock Center (BDSC). Two-way ANOVA, F_(12, 390)_ = 5.188, p< 0.0001 followed with Šídák’s multiple comparisons test. (B) Sleep duration is decreased in *inc^RNAi^*(P = 0.0098) and *CG5151^RNAi^* flies (P = 0.0307) on food and the knockdown of *fit* (P = 0.0374) or *inc* (P = 0.0465) in LHLK neurons inhibits the sleep loss induced by food deprivation. Two-way ANOVA, F_(13, 420)_ = 8.480, p< 0.0001 followed with Šídák’s multiple comparisons test. N = 16-20 per group. Data are mean ± SEM; *P < 0.05; **P < 0.01; ***P < 0.001.

**Figure Supplemental 2.**
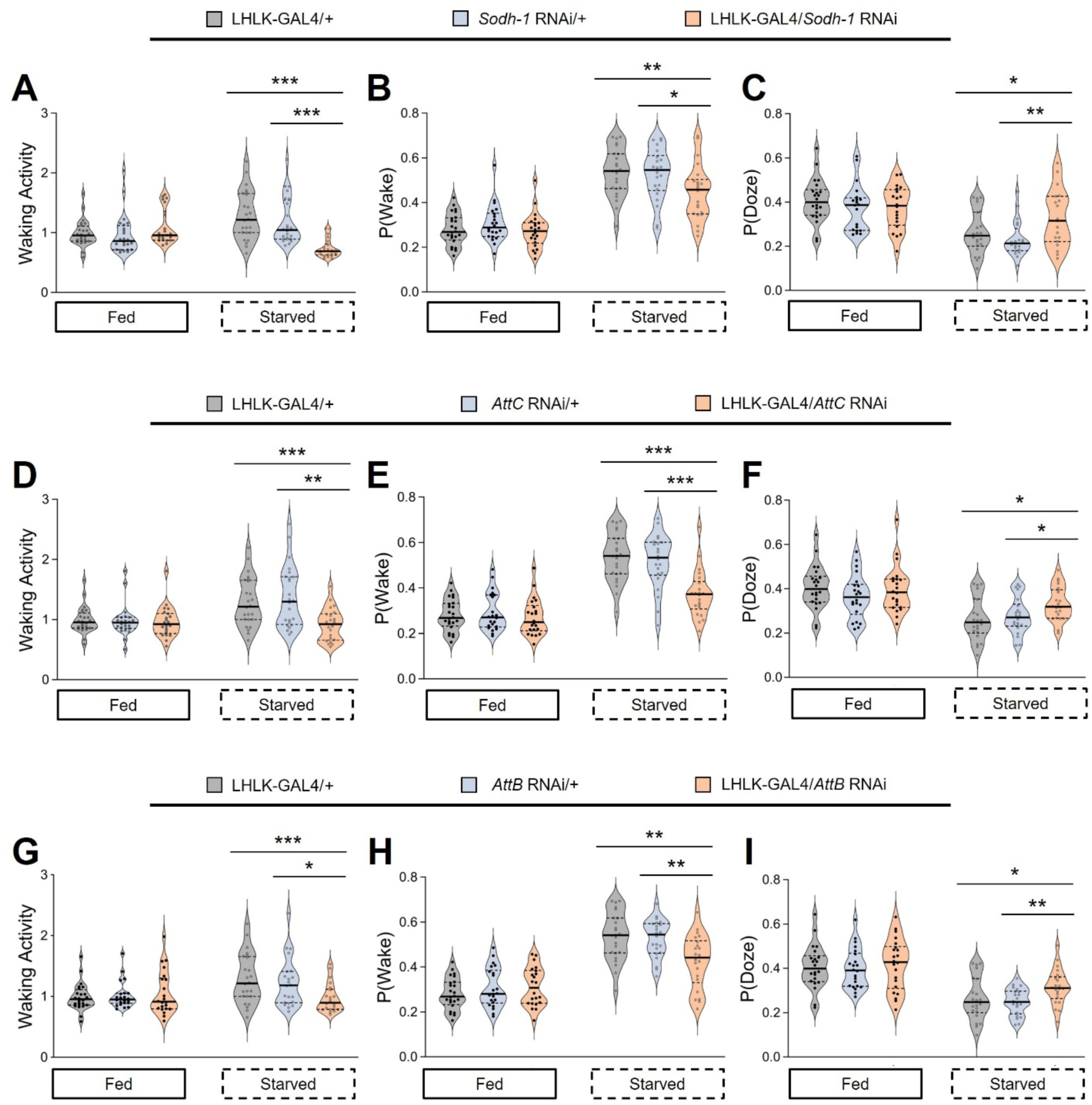
Characteristics of sleep and wakefulness modulated by knockdown of *Sodh1* or *Attacins* in LHLK neurons. **A**,**D**,**G**) Waking activity in mutants does not differ from two control groups (light gray, light blue) under fed conditions and is reduced compared to controls under starved conditions by knockdown of *Sodh1* (**A**, P < 0.0001), *AttC* (**D**, P = 0.0009) or *AttB* (**G**, P = 0.0028) in LHLK neurons respectively. **B**,**E**,**H**) The waking propensity p(Wake) of three different knockdown flies does not differ from controls under fed conditions, and is decreased in starved flies (P = 0.0117 for LHLK-GAL4>*Sodh1*^RNAi^ in **B**, P < 0.0001 for LHLK-GAL4>*AttC*^RNAi^ in **E**, and P = 0.0017 for LHLK-GAL4>*AttB*^RNAi^ flies in **H**). **C**, **F**, **I**) The sleep propensity p(Doze) is increased in knockdown flies, which targeting *Sodh1* (**C**, P = 0.0085), *AttC* (**F**, P = 0.0115), or *AttB* (**I**, P = 0.0486) with a driver for LHLK neurons, compared to controls on agar, but not on food. Kruskall-Wallis with Dunn’s post hoc. An unpaired t-test was used for comparisons between the LHLK knockdown group and a certain genetic background control respectively. *P < 0.05, **P < 0.01 and ***P < 0.001. Data are mean ± SEM.

**Table S6.**
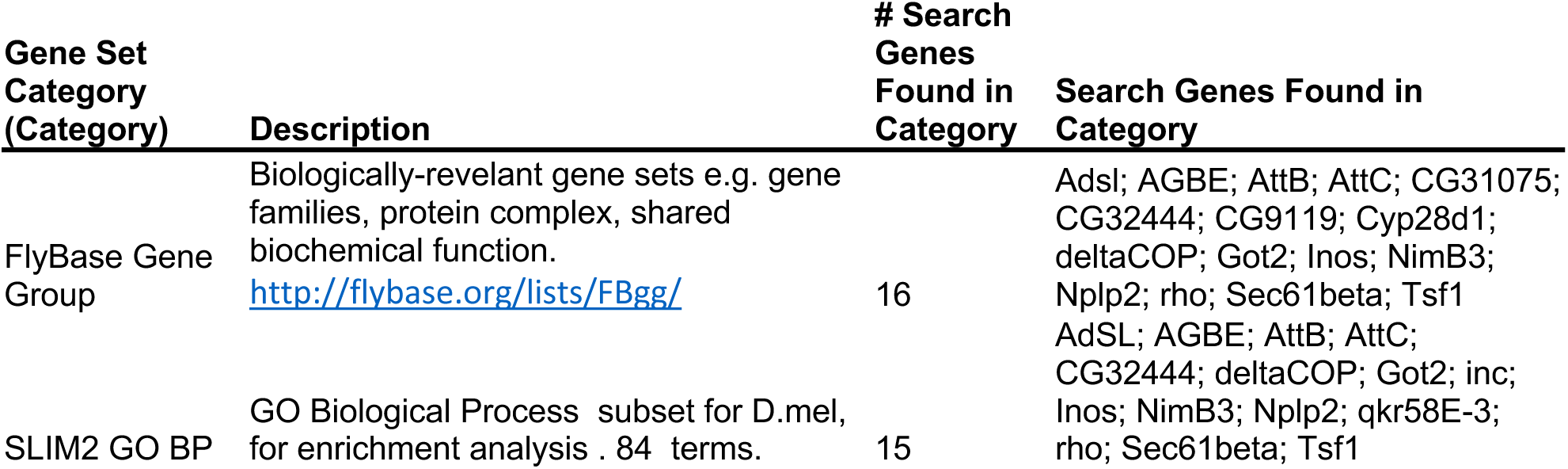
Two Gene Sets Selected for GO analysis

| Gene Set Category (Category) | Description | # Search Genes Found in Category | Search Genes Found in Category |
| --- | --- | --- | --- |
| FlyBase Gene Group | Biologically-relevant gene sets e.g. gene families, protein complex, shared biochemical function.<br>http://flybase.org/lists/FBgg/ | 16 | Adsl; AGBE; AttB; AttC; CG31075; CG32444; CG9119; Cyp28d1; deltaCOP; Got2; Inos; NimB3; Nplp2; rho; Sec61beta; Tsf1 |
| SLIM2 GO BP | GO Biological Process subset for D.mel, for enrichment analysis . 84 terms. | 15 | AdSL; AGBE; AttB; AttC; CG32444; deltaCOP; Got2; inc; Inos; NimB3; Nplp2; qkr58E-3; rho; Sec61beta; Tsf1 |

**Table S7.** GO analysis results

| Gene Set Category | Gene Set Name | Gene Set Size | Count Overlap Gene | log2 fold | P value | Overlapping Gene Symbols |
| --- | --- | --- | --- | --- | --- | --- |
| FlyBase Gene Group | ATTACINS | 4 | 2 | 8.185 | 1.694e-5 | AttC; AttB |
| FlyBase Gene Group | ANTIMICROBIAL PEPTIDES | 25 | 2 | 5.541 | 8.286e-4 | AttC; AttB |
| SLIM2 GO BP | carbohydrate metabolic process | 264 | 3 | 2.725 | 1.007e-2 | CG32444; Inos; AGBE |
| SLIM2 GO BP | immune response | 309 | 3 | 2.498 | 1.538e-2 | AttB; Nplp2; AttC |
| FlyBase Gene Group | ISOMERASES | 250 | 2 | 2.219 | 6.804e-2 | CG32444; Inos |
| SLIM2 GO BP | respiratory system development | 271 | 2 | 2.103 | 7.827e-2 | rho; deltaCOP |

## Acknowledgement

We thank Y. Aso and G. Rubin (Janelia Research Campus) for the GFP reporter; We thank W.W. Liao (Washington University in St. Louis) and Y. Jin (Cold Spring Harbor Laboratory) for technical advice in analysis of sequence data; R. Eifert (Cold Spring Harbor Laboratory) for making some machined apparatus, M. Brill and A. Crocker (Middlebury College) for the advice regarding single-cell harvesting. We also are grateful to J. Beshel, R. Keegan, Y.-H. Chang, and L. Talbot for helpful discussions. This work was supported by US National Institutes on Deafness and Other Communication Disorders [5R01DC013071-06 to J.D.]; DART NeuroScience LLC [to J.D.]; and US National Science Foundation XSEDE allocation [TG-IBN170003 and TG-IBN190002 to M.F.M.S.].

This work utilized the computational resources of the NSF CyVerse (www.cyverse.org, supported by DBI-0735191, DBI-1265383, and DBI-1743442) and Jetstream at IU/TACC (under XSEDE and supported by ACI-1548562). This work was supported by NINDS R01 NS131628 to ACK and JD.

---

|  |  |
| --- | --- |
| Fig 1 | Single-cell transcriptome analysis of LHLK neurons via Patch-Seq. |
| Fig 2 | Transcriptional analysis of LHLK neurons reveals differentially expressed genes |
| Fig 3 | Transcriptional reduction of Sodh1 or Attacins in LHLK neurons mitigates starvation-induced sleep suppression |
| Fig 4 | Behavioral role for genes downregulated in LHLK neurons following food deprivation. |
| Fig 5 | Starvation-induced sleep loss is reduced by knockdown of up-regulated or down-regulated candidate genes |
| Tab 1 | Preliminary sequencing hits |
| File S1 | CEL-Seq2 Protocol Adapted for GFP-labeled Neurons in flies |
| Fig S1 | Preliminary RNAi screening for genes up-regulated or down-regulated in LHLK neurons following 24 hours food deprivation. |
| Fig S2 | Characteristics of sleep and wakefulness modulated by knockdown of Sodh1 or Attacins in LHLK neurons |
| Tab S1 | sc-qRT-PCR_LHLK |
| Tab S2 | scRNASeq_41LHLK_count |
| Tab S3 | scRNASeq_41LHLK_TPM |
| Tab S4 | scRNASeq_18starved-23fed_2kVars.DESeq2_lib-state |
| Tab S5 | Two Gene Sets Selected for GO analysis |
| Tab S6 | GO analysis results |
| Tab S7 | Behavioral screening strains and results |

